# Weight loss increases adipose dendritic cells, non-classical antigen presenting proteins, and cytotoxic effector cells

**DOI:** 10.64898/2026.07.13.738291

**Authors:** W. Khan, M. Kapadia, E. Mancera, E.C. LaVoy, H. Wu, H.L. Caslin

**Affiliations:** Department of Health and Human Performance, University of Houston, Houston, TX, USA; Department of Medicine, Baylor College of Medicine, Houston, TX, USA

## Abstract

Weight cycling (i.e. cycles of weight gain, loss, and regain) is a growing health concern that has been shown to worsen risk of diabetes beyond that of obesity. Dendritic cells play a causal role in obesity-associated inflammation and metabolic disease. However, whether dendritic cells are impacted by weight cycling is not known. Here, we aimed to test the hypothesis that antigen presentation increases in dendritic cells with weight cycling. C57Bl/6J male mice were put on nutrient-matched low-fat or high-fat diets to elicit lean, weight gain, weight loss, and weight cycled groups. Adipose tissue immune cell populations and proteins related to signal 1, 2, and 3 of antigen presentation were analyzed by single cell-RNA sequencing and flow cytometry. Total adipose tissue dendritic cells, conventional type I and II dendritic cells, and monocyte-derived dendritic cells all increased with weight loss. Regarding major histocompatibility complex (MHC) protein expression, non-classical MHCIb proteins (Qa1, Qa2 and CD1d) were also highest in the weight loss group. Dendritic cells expressing the costimulatory molecule CD86 and the cytokine TNF were highest in the weight loss group. Finally, we assessed effector immune cell populations in the adipose tissue to understand the functional role of antigen presentation. CD8+ T cells, natural killer (NK) T cells, and NK cells also increased consistently with weight loss, but showed evidence of exhaustion. In sum, weight loss, but not weight cycling expands dendritic cells, increases signals for non-classical antigen presentation, and expands CD8+ T cells, NKT cells, and NK cells in the adipose tissue.

## Introduction

Obesity is a major risk factor for diabetes and cardiovascular disease^1^. One mechanism by which this can occur is through chronic inflammation^2,3^. Immune cells such as macrophages and T cells shift from a regulatory (“anti-inflammatory”) phenotype in lean adipose tissue to a more inflammatory phenotype following adipose expansion, which has been observed in both humans and mouse models^4–6^. These inflammatory populations secrete cytokines including IL-6 and TNF that impair insulin secretion and signaling and promote diabetes risk^7^. However, therapeutic interventions targeting inflammatory cytokines have not been clinically successful in diabetes prevention^8^, highlighting a gap in our understanding of obesity-induced inflammation.

Weight loss is an effective way to reduce obesity-related conditions including insulin resistance^9^. However, weight regain is common, and it is estimated that more than 30% of the population goes through cycles of weight gain, loss, and regain over their lifetime^10–12^. This repeated process of gaining, losing and regaining weight is known as weight cycling^13,14^. Notably, weight cycling worsens risk of diabetes, cardiovascular disease, and all-cause mortality more than obesity itself^14–17^. Importantly, our lab and others have observed that obesity-induced changes persist following weight loss and weight regain. For example, macrophages retain and even amplify inflammatory capacity following weight loss^18–21^. Moreover, upon weight regain, macrophages have higher basal cytokine production than in obesity, suggesting that weight cycling affects innate immunity^22,23^. Weight loss also drives clonal CD8+ T cell memory^20^, and blocking T cell memory formation through CD70 improves glucose tolerance after weight regain^24^, suggesting a role for adaptive immunity in weight cycling. However, any link between the two arms of the immune system during weight loss and weight cycling is not clear.

Antigen presenting cells link innate and adaptive immunity^25^. Dendritic cells, the primary antigen presenting cells, present endogenous or exogenous peptide antigens to T cells through major histocompatibility complex (MHC) I or II molecules following innate activation^26^. Notably, dendritic cells also play a causal role in obesity-associated inflammation and metabolic disease^27–30^. In mice and humans with obesity, adipose tissue dendritic cell populations positively correlate with increased insulin resistance^30,31^. In mice lacking dendritic cells, there is reduced weight gain and improved insulin tolerance following high fat diet^27,30^. Furthermore, injecting activated bone marrow-derived dendritic cells into mice on high fat diet increases adipose tissue inflammation^27,32^. However, any role for dendritic cells in weight cycling- associated disease has not been directly examined. In this study, we aimed to determine if adipose dendritic cells and antigen presentation-related proteins increase with weight loss and weight regain.

We found that some adipose dendritic cell populations increase with weight gain, but all populations increase further with weight loss. Additionally, non-classical MHCI molecules Qa1, Qa2, and CD1d, the costimulatory molecule CD86, and TNF expression increase consistently with weight loss. Finally, NK, NKT, and CD8+ T cells also increase with weight loss, suggesting that weight loss, but not weight regain, increases dendritic cell antigen presentation and many adaptive immune cells.

## Methods

### Mice

All experiments were completed at the University of Houston in compliance with the University of Houston Institutional Animal Care and Use Committee (PROTO202300026). The University of Houston is accredited by the Association for Assessment and Accreditation of Laboratory Animal Care International.

Male C57BL/6J mice were purchased from Jackson Labs (#000664) at 7 weeks of age. At 8-9 weeks of age, mice were placed on cycles of a high-fat diet (HFD; 60% fat, Research Diets #D12492) or a low-fat diet (LFD; 10% fat, Research Diets #D12450J) for 27-29 weeks as previously published and in **Fig. 1A** ^20,33^. During the study period, mice had unrestricted access to water and food, and body weight was measured weekly.

**Figure 1:**
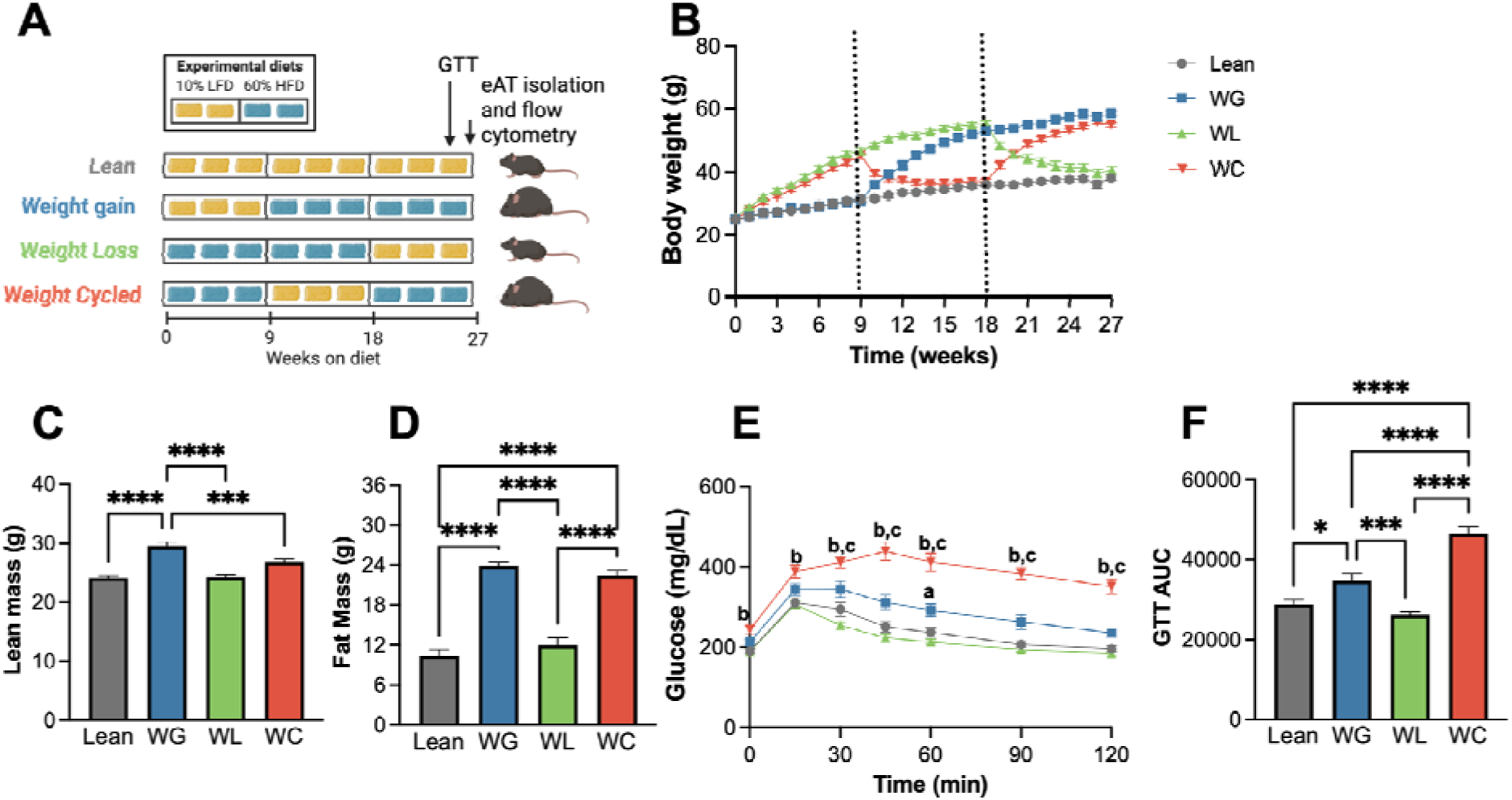
Diet-induced weight cycling worsens glucose tolerance. A) Schematic of model. B) Body weight over time in lean, weight gain (WG), weight loss (WL) and weight cycled (WC) mice diet change upon 9 weeks (shown as dotted lines). C) Lean mass, D) fat mass, and E) Blood glucose over 2 hours during an intraperitoneal glucose tolerance test (1.5 g dextrose/kg lean mass) at end of study. Lowercase letters a = Lean vs WG, b = Lean vs WC, c = WG vs WC, indicate statistically significant differences (p < 0.05) among groups at a specific time point F) Area Under the Curve (AUC) of glucose tolerance test. * p<0.05; ***p<0.0005; ****p<0.0001.

### Body composition

Lean and fat mass were measured with magnetic resonance imaging (EchoMRI - 100 Body Composition Analyzer) at 26 weeks in all mice.

### Glucose tolerance testing

After body composition testing, an intraperitoneal glucose tolerance test (ipGTT) was performed in mice to measure glucose tolerance. After a 5-hour fasting period, baseline blood glucose levels were measured. Next, mice received an intraperitoneal injection of dextrose at a dose of 1.5 g/kg lean mass, and blood glucose levels were monitored by tail nick at 15, 30, 45, 60, 90, and 120 minutes post-injection using hand-held glucometer (CVS Advanced Glucose Meter).

### Immune cell isolation from adipose tissue

After 27-29 weeks on diet, mice were euthanized by carbon dioxide exposure followed by cervical dislocation and subsequently perfused with 20 mL of PBS through the left ventricle. The epididymal adipose fat pads were collected and weighed. Adipose tissue was then minced and digested in 2 mg/mL type IV collagenase (Worthington, #LS004176) for 30 minutes at 37°C. Following digestion, the suspension was vortexed and passed through a 100-µm filter with cold PBS, and the adipocytes were removed by centrifugation. The red blood cells were then lysed with ACK lysis buffer (Quality Biological, #118-156-101), and the stromal vascular fraction was subsequently filtered through a 35-µm strainer for immune cell analysis.

### Flow cytometry

The stromal vascular fraction was used to measure the frequency of adipose tissue dendritic cells and the expression of antigen presenting proteins, co-stimulatory molecules, and intracellular cytokines. Zombie Violet viability stain (BioLegend, 423114) was added 10 min in protein free buffer followed by FcBlock (BioLegend, 156604) for 10 min. Surface staining was then completed for 30 minutes at 4°C with the following fluorescently labeled antibodies: BV510– CD45 (BioLegend, 103138), FITC–CD11c (BioLegend, 117306), PerCP–CD11b (BioLegend, 101228), PE-Cy7–CD301b (BioLegend, 146807), APC–CD64 (BioLegend, 161006), APC-Cy7– XCR1 (BioLegend, 148223), PE–H2 MHC I (BioLegend, 125505), PE–H2-Q7/Qa2 (BioLegend, 121715), PE-Cy7–I-A/I-E (BioLegend, 107629), APC Fire–H2Db (BioLegend, 111519), PE–CD1d (BioLegend, 123509), PerCP-Cy5.5-H2K-b (BioLegend, 116515), PE-Qa1b (BD Pharmingen), PE-Cy7–CD86 (BioLegend, 159207), APC-Cy7–CD40 (BioLegend, 124637), FITC-NK1.1 (BioLegend, 108705), APC-Cy7-CD3 (BioLegend, 100211), PE-Cy7-CD49a (BioLegend, 142607), and APC-CD8 (BioLegend, 100711). All cells were fixed before flow cytometry with fixation buffer (from the FoxP3 Transcription Factor Staining Buffer Set (eBioscience/ Thermo, 00-5523-00). For intracellular cytokine staining, the full FoxP3 Transcription Factor Staining Buffer Set was used to fix and permeabilize the cells. Intracellular staining was completed overnight at 4°C with the following fluorescently labeled antibodies: PE- TNF (BioLegend, 506306). Fluorescent minus one (FMO) controls were used to assist with gating. Cells were run on a Miltenyi Quant10 flow cytometer and data was analyzed on Flow Jo software (version 10). The gating strategy used to define dendritic cell subsets and the expression of MHCs, costimulatory molecules, and TNF is shown in Supplementary Figure S1. The gating strategy used to define NK, NKT and CD8+ T cells is shown in Supplementary Figure S2.

### Single cell RNA sequencing analysis

To further examine dendritic cell subpopulations, we completed additional analysis of a previously published single cell sequencing dataset with epididymal adipose tissue stromal vascular fraction samples from the same weight cycling model (GEO accession number <u>GSE182233</u>^20^). Violin plots were generated with R version 4.5.2 and Seurat version 5.4.0^34^.

### Statistical analysis

Two independent experiments were completed. Mouse phenotype data and total dendritic cell data was combined for analysis. Qa2 and CD1d are representative of the two independent experiments. All other population measures were measured once, but confirmed with sequencing.

For all murine and flow cytometry data, statistical analysis was conducted using GraphPad Prism version 10.5.0 software. Outlier testing and normality testing were run on all data. For data with a normal distribution, a one-way analysis of variances (ANOVA) was used. When the F statistic was significant, a Tukey multiple comparison test was used to identify between group differences. For data that did not pass normality testing, Kruskal-Wallace non- parametric tests were used. Post-hoc testing was performed with a Dunn’s multiple comparison test when appropriate. All data is presented as mean ± standard error (SEM) and significance was set at p-value < 0.05.

## Results

### Weight cycling worsens glucose tolerance

To determine the impact of weight loss and weight cycling on dendritic cell populations, we used cycles of high-and low-fat diet feeding to generate lean, weight gain, weight loss, and weight cycled mice (Fig 1A) and as previously published^20,33^. At the end of the study period, lean and weight loss groups had similar total body mass, lean mass, and fat mass (Fig 1B-D). Similarly, the weight gain and weight cycled groups had similar total body mass and fat mass at study end, although the lean mass was slightly higher in the weight gain group (Fig 1B-D). Importantly, we showed via ipGTT that glucose tolerance was significantly impaired in the weight gain group vs. lean group, and glucose tolerance was further impaired in weight cycling vs. weight gain group (Fig 1E&F), as seen in the previous cohorts^20,33,35^. These data support the use of our mouse model to replicate weight cycling-accelerated diabetes risk.

### Weight loss increases dendritic cell subpopulations in adipose tissue

To investigate adipose tissue dendritic cells during weight cycling, we first assessed population changes by flow cytometry. As a percent of CD45+ cells, adipose tissue dendritic cells (CD45+CD11c+ CD64-)^31^ significantly increased with weight loss compared to lean, obese, and weight cycled mice (Fig 2A). The total number of dendritic cells per gram of adipose tissue also significantly increased in the weight gain, weight loss, and weight cycled groups compared to lean, with more dendritic cells in the weight loss vs. weight cycled group. We then examined dendritic cells further by subset. The conventional dendritic cell (cDC)1 population (XCR1+)^36^ significantly increased in number, but not percentage, with weight loss and weight cycling compared to the lean group (Fig 2B). The cDC2 population (XCR1-CD301b+CD11b+)^37^ increased significantly by percent of CD45+ cells and number with weight loss compared to lean and weight cycled mice (Fig 2C). Last, the moDC population (XCR1-CD11b+CD301-)^37^ significantly increased in percent of CD45+ and number of cells upon weight gain and weight loss compared to lean and weight cycled mice (Fig 2D). Together, these data suggest a general trend in which weight gain increases dendritic cell populations, which are further increased with weight loss and decrease with weight regain. Importantly, cDC2 and moDC increase by percent with weight loss and there is a consistent and significant increase in cDC1, cDC2, and moDCs by number with weight loss.

**Figure 2:**
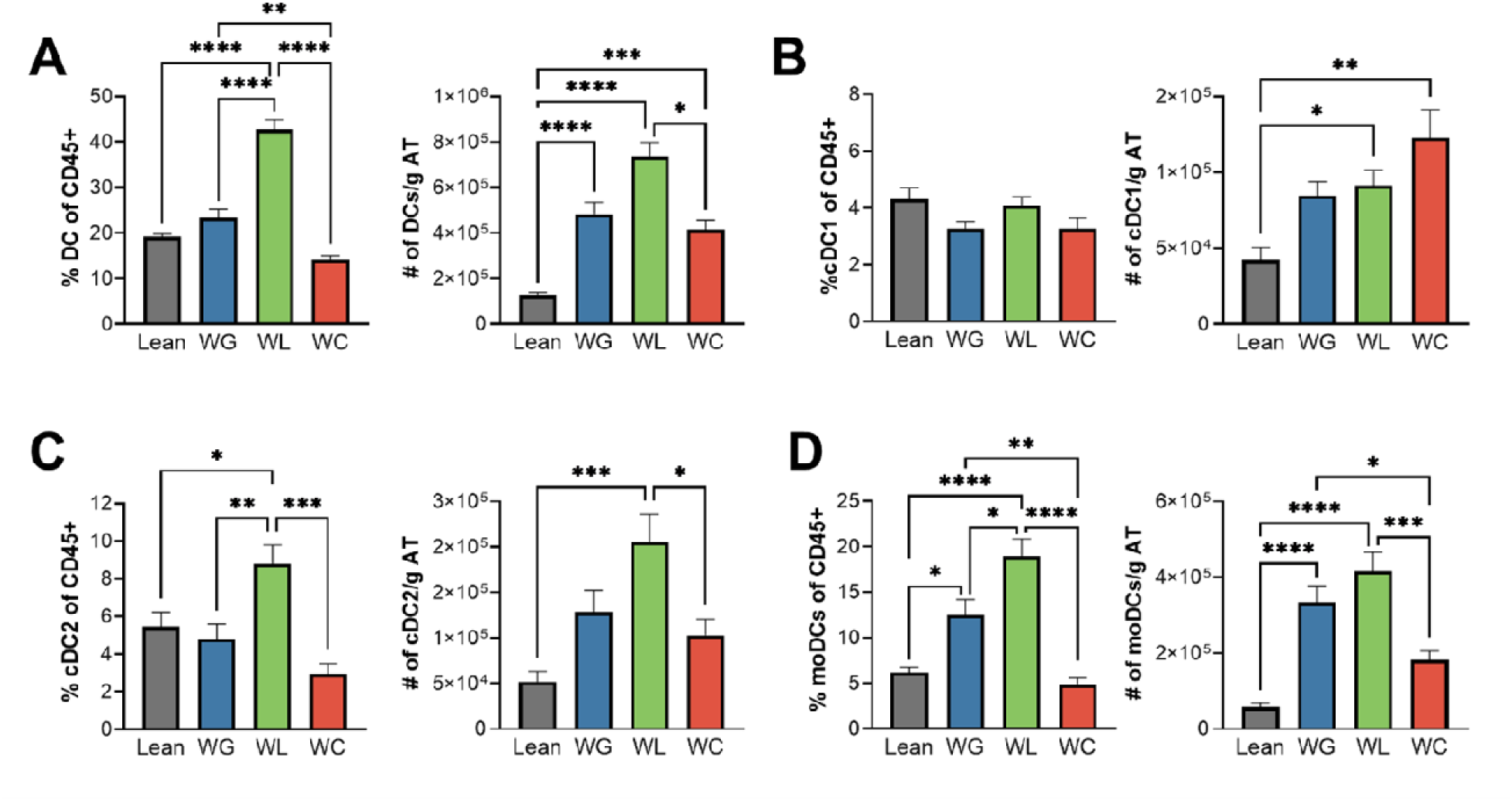
Weight loss increases cDC2 and moDCs in adipose tissue (AT). From left to right the percentage and absolute number per gram of epididymal adipose tissue for A) total dendritic cells (DCs; CD45+CD11c+CD64-), B) conventional DC1 (cDC1; DCs with XCR1+), C) conventional DC2 (cDC2; DCs with CD11b+CD301b+), and D) monocyte-derived DCs (moDCs; CD11b+CD301b-). * p<0.05; ** p<0.001; ***p<0.0005; ****p<0.0001.

### Weight loss increases the frequency and expression of Qa1+, Qa2+, and CD1d+ dendritic cells

Next, we analyzed the protein expression of MHCI and MHCII molecules across all (CD11c+CD64-) dendritic cells as an indication of their function for antigen presentation. There was a significant decrease in the percent of cells expressing the MHCII protein IA/IE upon weight gain and weight loss mice compared to lean, which was recovered with weight cycling (Fig 3A). However, among the cells positive for IA/IE, there was no change in IA/IE expression by median fluorescent intensity (MFI). As dendritic cells are often defined by MHCII, we ran new analysis on previously published single cell RNA-sequencing data^20^. We found that MHCII was still transcribed at the gene level in cDC1, cDC2, and moDC clusters (Fig S3), suggesting the influx of “pre-DC” or immature dendritic cells with little or no MHC expression on the surface^38–40^. Moreover, in our initial dataset, there were clusters of each sub-type with and without *Ccr7*, which can distinguish mature DCs^20^, suggesting the presence of both mature and immature dendritic cells in the adipose tissue. We also observed a general trend towards a decrease in percent of the classical MHCI protein H2Db with weight gain and weight loss, that was recovered with weight cycling (Fig 3B), however, there was no statistical difference between lean, weight gain, and weight loss cells, and there was no change in the MFI on positive cells. The percent of dendritic cells expressing another classical MHCI protein (H2Kb) significantly increased upon weight loss compared to the lean, weight gain, and weight cycled mice (Fig 3C). While this may not seem physiologically significant (proportions increased from 98-99%), the amount per cell also increased in the weight loss group (as observed by protein MFI (Fig3C) and gene expression (Fig S3)).

**Figure 3.**
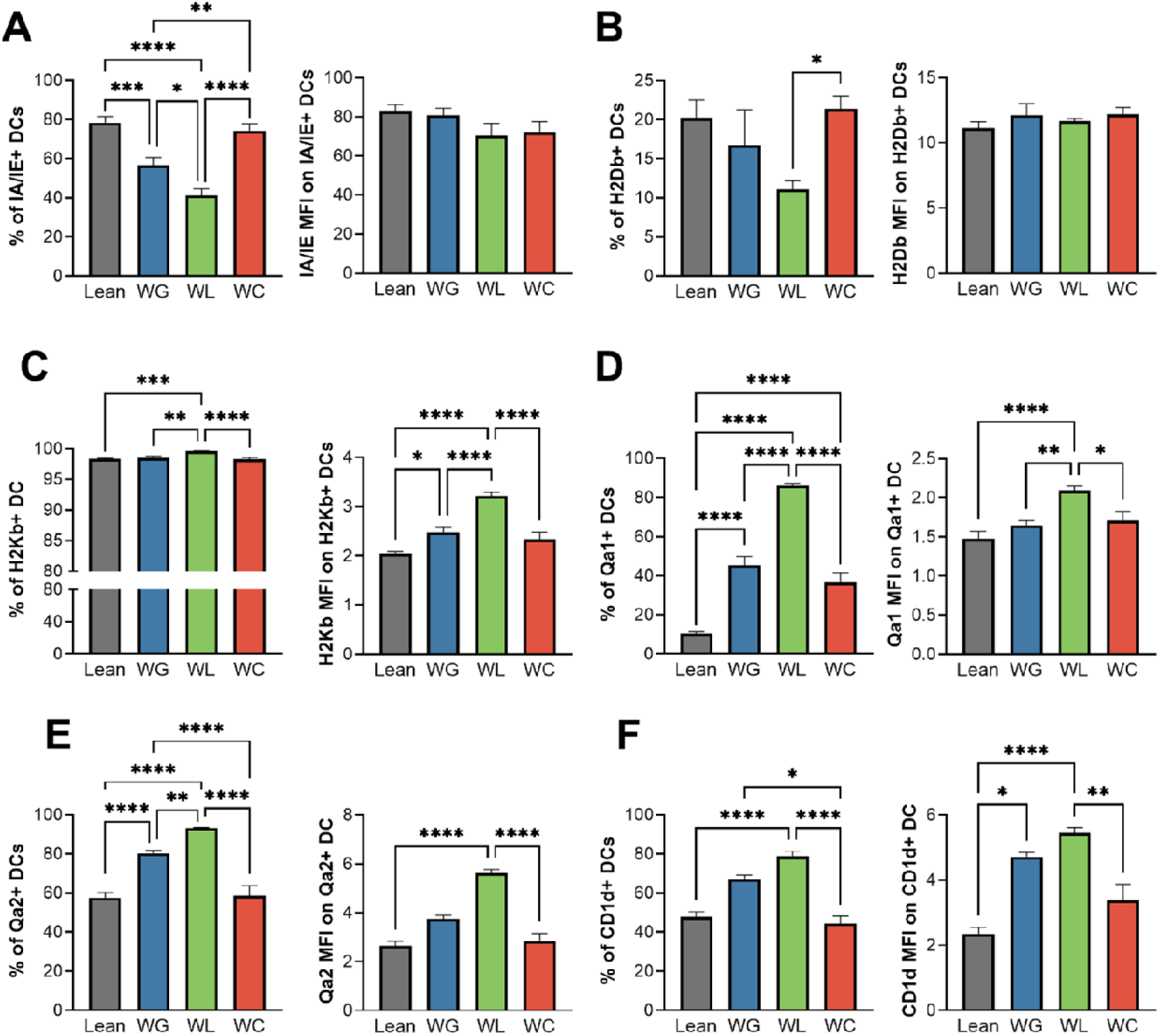
Weight loss increases the frequency and expression of the MHC molecules H2Kb+, Qa1+, Qa2+ and CD1d+ DCs. From left to right the percent and MFI (median fluorescent intensity) of A) IA/IE+, B) H2Db+, C) H2Kb+, D) Qa1+, E) Qa2+, and F) CD1d+ on total DCs (CD45+CD11c+CD64-). * p<0.05; ** p<0.001; ***p<0.0005; ****p<0.0001

Among the non-classical MHCI proteins, there was a significant increase in dendritic cells expressing Qa1 and Qa2 upon weight gain compared to lean and an even greater increase during weight loss (Fig 3D &E). The %Qa1+ dendritic cells remained elevated from lean in the weight cycled group, despite a reduction from weight loss, while the %Qa2+ cells returned to lean levels. Among those positive for Qa1 and Qa2, there was also an increase in the MFI of Qa1 and Qa2 in the weight loss compared to the lean and weight cycled groups, suggesting more cells expressing these MHC molecules and more molecules per cell following weight loss. In our sequencing data, the expression of *H2-Q6* and *H2-Q7* (Qa2) also increased with weight gain and had a further increase with weight loss in cDC1 and cDC2, and *H2-T23* (Qa1) increased in moDCs with weight loss, consistent with our flow cytometry data (Fig S3). We also observed a significant increase in the dendritic cells expressing the non-classical MHCI-related glycoprotein CD1d with weight loss compared to lean and weight cycled mice and an increase from lean in the MFI on CD1d+ cells with weight gain and weight loss (Fig 3F). Of note, we could not detect *Cd1d1* in the sequencing dataset (Fig S3). Together, these data suggest that weight gain and more so, subsequent weight loss, drives a consistent increase in non-classical MHCI proteins and likely antigen presentation.

Given that macrophages also function as professional antigen-presenting cells and are highly abundant in adipose tissue, we also examined the inflammatory macrophage population (CD11c+ CD64+) and their MHC expression in our flow cytometry data. The percent and total number of inflammatory macrophages significantly increased with weight gain and weight cycling (Fig S4A). Weight loss reduced inflammatory macrophages compared to the weight gain and weight cycled groups, but this population remained elevated compared to lean mice. While there was significantly less IA/IE per cell with weight gain and weight cycling, the macrophage population expressing IA/IE protein remained unchanged (Fig S4B), and the percent of macrophages expressing H2Db showed a slight increase with weight loss with no change in MFI (Fig S4C). Both results were different from dendritic cells. Interestingly, like the dendritic cell population, there was a significant increase in both percent and MFI of H2Kb+, Qa1+, and Qa2+ macrophages with weight loss (Fig S4D-F). However, unlike dendritic cells, the percent of CD1d+ macrophages decreased with weight gain and weight cycling with a significant decrease in the expression per cell in weight gain, weight loss, and weight cycled mice (Fig S4G). In sum, weight loss specifically increases classical MHCI H2Kb and non-classical MHCI Qa1 and Qa2 on both macrophages and dendritic cells. Additionally, dendritic cell expression of CD1d increases with weight loss. These results highlight differential regulation of MHC proteins in different cell populations and subsets with weight gain, weight loss, and weight cycling.

### Weight loss increases the costimulatory marker CD86 on dendritic cells

Antigen presentation via both MHCII and MHCI molecules is considered “signal 1” for T cell activation and is supported by the expression of co-stimulatory molecules, considered “signal 2”^41^. As we observed a significant increase in protein expression of many antigen presenting molecules, we next assessed the protein expression of the co-stimulatory molecules CD86 and CD40. Consistent with the observed increases in MHC molecules with weight loss, there was a significant increase in the percent of CD86+ dendritic cells in weight loss compared to weight gain and weight cycled mice and there was an increase in the MFI of CD86 on positive cells in the weight loss compared to lean and weight cycled mice (Fig 4A). However, while there was a trend towards an increase in both the percent and MFI of CD40+ dendritic cells with weight loss, it was not statistically significant (Fig 4B).

**Figure 4:**
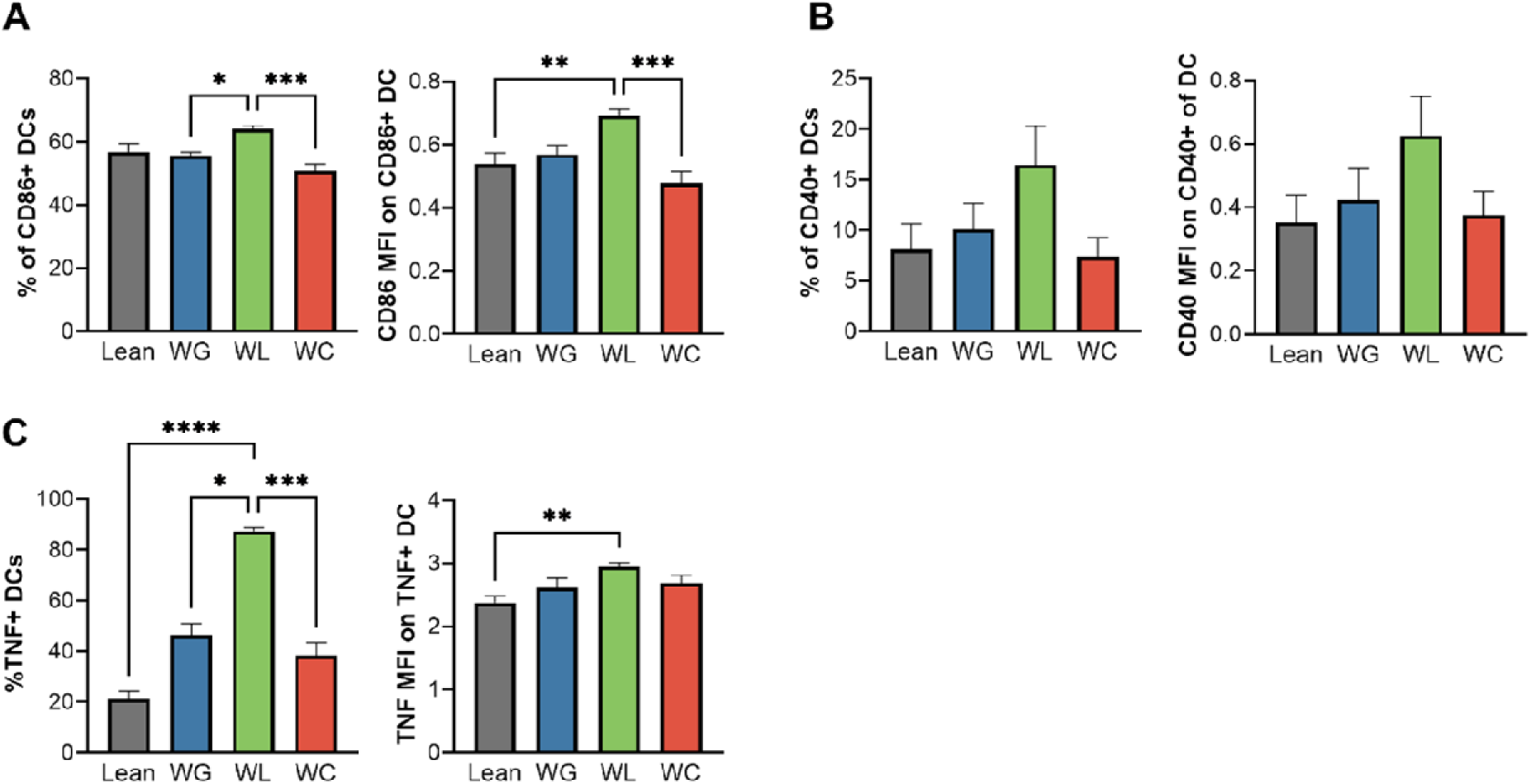
Weight loss increases the costimulatory marker CD86 and TNF cytokine production in DCs. From left to right the percent and MFI (median fluorescent intensity) of A) CD86, B) CD40, and C) TNF of total DCs (CD45+CD11c+ CD64-). * p<0.05; ** p<0.001; * **p<0.0005; ****p<0.0001.

Like above, we wanted to see how these and other costimulatory molecules change across subpopulations of dendritic cells. Analysis of our previously published single cell RNA-sequencing data showed that each subpopulation had unique expression of one or more costimulatory molecules (Fig S5). Interestingly, the expression of *Cd86* increased with weight gain and remained elevated in all subpopulations with weight loss. Only in moDC did *Cd86* further increase with weight loss (Fig S5C). Additionally, *Cd40* was only expressed in cDC2, showing an increase with weight gain and weight loss (Fig S5B). The expression of *Tnfsf9* (the gene for 4-1BBL) also increased with weight gain in cDC2 and moDC and even further increased with weight loss in cDC2, suggesting another potential mediator of interest (Fig S5B- C). Together, our data suggest that like MHC molecules, there is an increase in the expression of CD86 on dendritic cells with weight loss, which may be at least partly driven by cDC2. CD40 and 4-1BBL may also increase on cDC2 with weight loss, although further flow cytometric phenotyping is required to identify changes at the protein level.

Given the increased expression of MHC (signal 1) and costimulatory molecules (signal 2) on dendritic cells with weight loss, we next assessed inflammatory cytokine expression (signal 3), focusing on basal tumor necrosis factor (TNF) expression^42^. The percent of TNF-producing dendritic cells increased in the weight loss mice compared to all other groups and MFI on TNF positive dendritic cells increased with weight loss compared to lean controls (Fig 4C). Like before, we also looked back at the single cell sequencing data to understand how TNF and other inflammatory cytokines change across subpopulations (Fig S6). In the cDC2 and moDC population, the expression of *Tnf* increased with weight gain, decreased slightly with weight loss (yet remained elevated compared to lean), and then increased most with weight cycling (Fig S6B&C), suggesting modulation at the translational or post-translational levels or gene dropout with sequencing. In cDC1, *Il18* was expressed in lean and weight cycled cells (Fig S6A), and in moDC, *Il6* expression decreased upon weight gain, but increased again with weight loss and weight cycling, yet did not increase to levels observed in lean samples (Fig S6C). Thus, while there are subset specific costimulatory molecule and inflammatory cytokine profiles, and slight differences in gene expression and protein expression, dendritic cells generally increase signal 1 (H2Kb, Qa1 and Qa2), signal 2 (at least CD86), and signal 3 (at least TNF) with weight loss.

### Weight loss increases CD8+ T cells, NKT cells, and NK cells

Antigen presentation and co-stimulation ultimately modulate cytotoxic effector immune cell populations. H2Kb, Qa1, and Qa2 can present protein antigens to CD8+ T cells and CD1d presents lipid antigens to NKT cells^43–48^. Qa1 and 2 can also present to NK cells^48,49^. Therefore, we also looked at the innate and adaptive cytotoxic effector cells in the adipose tissue. Compared to lean mice, there was a significant increase in the number of CD8+ T cells and NKT cells with weight gain, weight loss, and weight cycling, which was seemingly highest in the weight loss group (Fig 5A and B). The proportion of CD8+ T cells trended up with weight loss and NKT proportion significantly increased in weight loss compared to lean and weight gain. Additionally, the weight loss mice had the greatest proportion and number of NK cells compared to all other groups (Fig 5C). These data were supported by the single cell sequencing data, where weight loss specifically drives effector memory CD8+ T cells, NKT, and NK cells (Fig S7).

**Figure 5:**
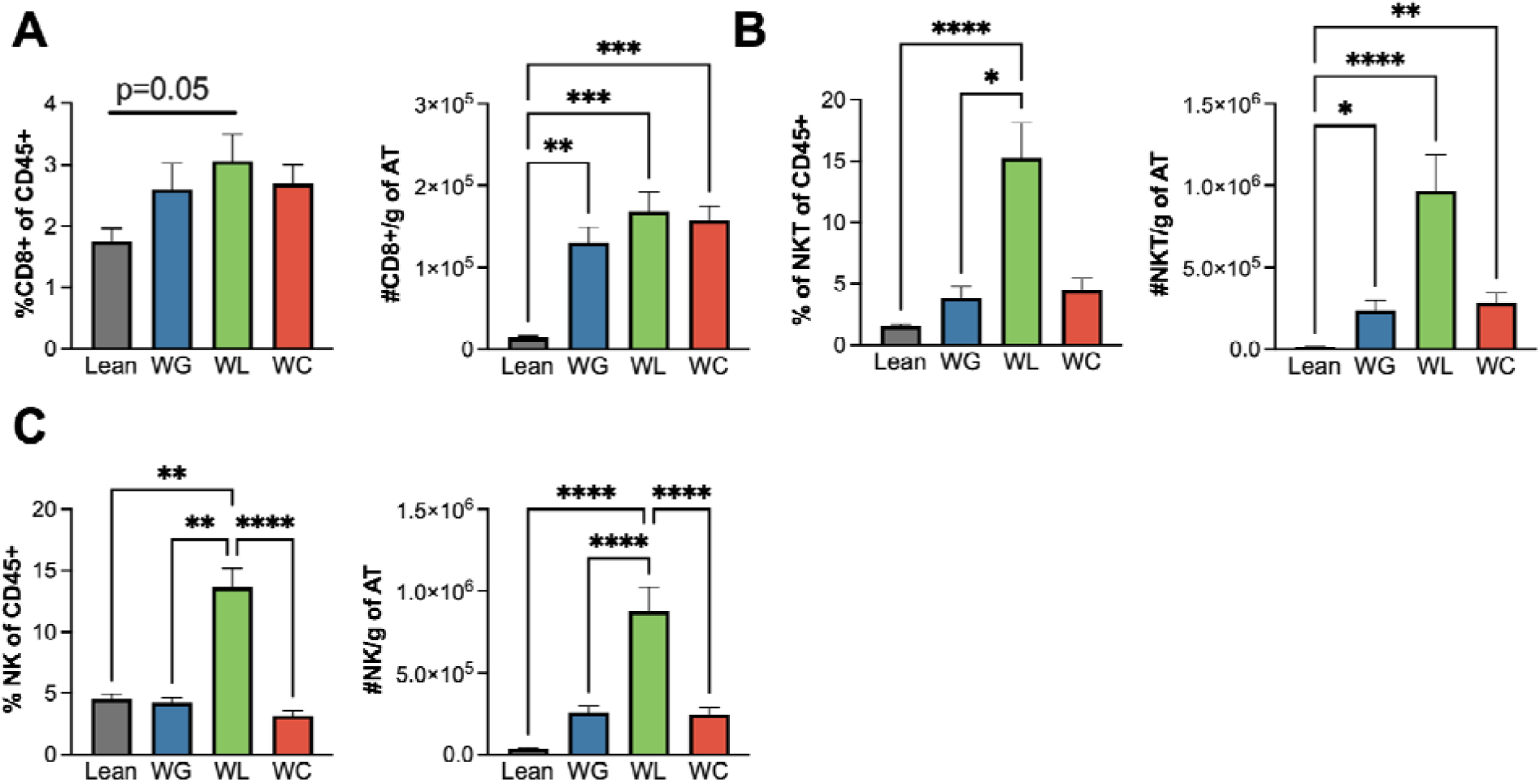
Weight loss increases CD8+ T, NKT, and NK cells in adipose tissue (AT). From left to right the percentage and absolute number per gram of epididymal adipose tissue for A) CD8+ T cells (CD3+NK1.1-CD8+), B) natural killer T cells (NKT; CD3+NK1.1+), C) natural killer cells (NK; CD3-NK1.1+). *p<0.05; ** p<0.001; ***p<0.0005; ****p<0.0001.

Weight gain has been shown to increase CD8+ T cell exhaustion^50^, and we have previously shown weight loss to further drive CD8+ T cell exhaustion^20^. Here, we find that NK cells also show reduced activation marker expression (*Itga4, Klrg1, Klra8*) and increased expression of *Tigit*, considered a marker of exhaustion *(Fig S8A and B*). As the inhibitory receptors *Klrc1, Klra3, and Klra9* were also reduced with weight loss (Fig S8C), these results suggest that non-classical antigen presentation may drive chronic antigen stimulation or activation, expansion, and exhaustion via mature dendritic cells vs. tolerance via immature cell populations^51^. Collectively, these results suggest that weight loss following a bout of weight gain increases adipose dendritic cells, three signals for non-classical MHC1b antigen presentation, and CD8+ T cell, NKT, and NK populations with evidence of exhaustion.

## Discussion

Weight cycling (i.e. previous weight gain, loss, and regain) exacerbates the risk of metabolic disease even greater than obesity itself^14,52^. It is well known that dendritic cells in adipose tissue directly contribute to inflammation and diabetes risk in obesity^27,28,30,31,53^, however, their role in disease risk associated with weight cycling is not clear. We hypothesized that antigen presentation would increase with weight cycling. Surprisingly, we found that weight loss following an initial bout of weight gain is the metabolic state that specifically increases adipose dendritic cell populations and non-classical MHCI antigen presentation. While we seemingly found evidence of “pre” or immature dendritic cell infiltration, we also found an increase in non-classical signal 1, 2, and 3 and cytotoxic effector immune cell expansion with evidence of exhaustion. Together, these data suggest that weight loss drives dendritic cell recruitment and maturation, antigenic activation, and immune dysregulation that may have further tissue implications upon weight regain. These results also prompt many new questions pertaining to the role of cytotoxic effector cells in the adipose tissue.

In this current investigation, we observed that while some dendritic cell populations in the adipose increase with weight gain, all populations increase by cell number following weight loss. Additionally, cDC2 and moDCs increased by percent with weight loss and cDC1 increased even further by cell number with weight regain. This work supports previous studies showing dendritic cells accumulate in adipose tissue in humans and mice with obesity^27,30,31,54,55^, but is one of the first to show broadly that dendritic cell populations further increase in proportion and number following weight loss and selectively in weight regain. In previous work, we showed that “activated” (*Ccr7*+) cDC1 and cDC2 increase during weight loss by single cell RNA-sequencing^20^ and another interesting study showed that just 36-hour fasting increased cDC1, but not cDC2, in adipose tissue^56^, suggesting some conserved effects of weight loss but also some unique regulation possibly due to the magnitude of weight or the impact of prior weight gain. As both cDC1 and cDC2 have homeostatic tolerogenic or suppressive functions in lean adipose tissue and both seemingly promote inflammation and worsen glucose tolerance following high fat diet feeding^57,58^, we are interested to understand how the increase in dendritic cells may affect cell function with weight loss.

To further understand how weight loss and weight regain may impact antigen presentation, we also measured MHC expression. Our data showed a decrease in the MHCII molecule H2-IA/IE with weight gain and loss, which was surprising as many published studies have shown a role for MHCII antigen presentation in the development of adipose inflammation and metabolic disease^59–61^. However, most of that published work suggests a regulatory role for adipocyte and macrophage MHCII and crosstalk with CD4+ T cells. Moreover, dendritic cells are typically defined by the expression of MHCII, and thus, we do not expect dendritic cells to lose MHCII expression with weight gain and loss. It is plausible that weight loss can drive “pre” or immature dendritic cell recruitment to the tissue, as these cells express little or no MHCII on the surface^59–62^. Additional evidence for this comes from our prior single cell sequencing work where we showed there are both “activated” (or mature^63^) *Ccr7*+ dendritic cells and *Ccr7*- negative dendritic cells go up with weight loss^20^. Weight loss is characterized by substantial lipolysis and not just lipid release, but also tissue remodeling and immune activation^64^.

Additionally, dendritic cells are relatively short-lived, with lifespans ranging from days to weeks depending on tissue and subset, and thus, are continuously replenished by circulating precursors^65,66^. Therefore, it is logical that weight loss over 9 weeks may increase dendritic cell recruitment, antigen exposure, and maturation and non-classical antigen presentation once in the tissue. Moreover, it is possible some of our CD45+CD11c+CD64- cells were B cells^67^ and non-classical monocytes^68^, which also increase with weight loss (Fig S9). It is also notable that non-classical monocytes also have low expression of MHCII and terminally differentiated plasma cells lose MHCII expression^69,70^, which may also help explain our MHCII data and have an effect on the adipose tissue. However, we expect that much of our non-classical MHCIb data is primarily due to dendritic cells. The observed increase in Qa1 appears specific to the dendritic cell populations, as *H2-T23* (Qa1) goes down by sequencing in B cells, plasma cells, and non-classical monocytes (Fig S9). Moreover, while *H2-Q6* and *H2-Q7 (*Qa-2) increase with weight loss in plasma and non-classical monocytes, these populations are substantially lower in abundance than the dendritic cells (Fig S3 and S9).

Our primary finding was an increase in classical MHCI H2-Kb and the non-classical MHCIb molecules Qa1, Qa2 and CD1d on dendritic cells with weight loss supported by sequencing and flow cytometry data. These observations indicate MHCIb may be the dominant antigen presentation pathway in dendritic cells upon weight gain and weight loss. Interestingly, we also found Qa1 and Qa2 increased on adipose macrophages with weight loss, suggesting again some conserved and some unique pathways regulated upon weight loss. We have not found any previous work on Qa1 or Qa2 in the adipose tissue, and like MHCII, prior studies on CD1d have primarily looked at macrophages and adipocytes^71–74^. Qa1 and Qa2 present protein antigens to CD8+ T cells and NK cells and CD1d presents lipid antigens to iNKT cells^43,44,46–49^. These antigens are considered more limited or restricted than the diverse range of antigens binding to classical MHCI and II^48,75^, however, the identity of any antigens presented with weight loss are not known. Moreover, many studies in development, homeostasis, cancer, and viral infections suggest that non-classical MHCI molecules are upregulated and suppress cytotoxic function^76–80^, however, there is also evidence that MHCIb-restricted CD8+ T cells can be clonally activated for enhanced viral and tumor control^43,81,82^. Our data suggest that non-classical MHCIb presentation also drives the activation and expansion of CD8+ T cells and NK cells.

We found that in addition to an increase in expression of non-classical MHCIb molecules (signal 1), there was an increase in CD86 (signal 2) along with an increase in TNF (one cytokine involved in signal 3) upon weight loss. There was also an increase in *Cd40* expression in cDC2 selectively and an increase in *Tnfsf9* (4-1BBL) specifically on cDC2 with the highest expression upon weight regain. CD86+ dendritic cells have been shown to negatively correlate with regulatory T cells^55^, however, dual B7 (CD80/CD86) depletion worsens insulin resistance despite reduced visceral adiposity under both normal and high-fat diets in mice^83^. Moreover, 4- 1BBL has been previously associated with aggravating adipose tissue inflammation and glucose intolerance in mice on high fat diet ^84^. Thus, future studies will need to more directly interrogate how specific MHC and co-stimulatory molecule expression mediates dendritic cell subtypes in weight gain, weight loss, and weight regain.

Importantly, we also found that weight loss increased CD8+ T cell, NKT, and NK populations, which supports protein expression data to suggest antigen presentation is functionally increased and induces effector expansion with weight loss. Previously published work supports our findings that these populations increase with obesity in adipose tissue^85–87^, yet this current study is one of the first to demonstrate that the number and proportion of these cell populations are further increased with weight loss. Prior work also suggests CD8+ T cells and NK cell promote inflammation and glucose intolerance in mice on high fat diet^88,89^, while invariant (i)NKT cells protect against diet induced obesity and glucose intolerance^90–93^.

However, the role of these cells during weight loss and any subsequent impact on weight regain is not known. We often consider cytotoxicity the primary function of these cells. iNKT cytotoxic function helps promote the turnover of adipocytes in a lean state^91^, but CD8+ T cells also expand within days of high fat diet feeding, secrete inflammatory cytokines, recruit macrophages, and worsen glucose tolerance in obesity^85,88,94^. The evidence of exhaustion in both CD8+ T cells and NK cells suggests chronic antigen stimulation^95–97^, which could act to restrain cytotoxic and inflammatory function and may serve to limit chronic inflammation. However, many recent studies also suggest exhaustion can also be an alternative activation state in which granzyme K production promotes inflammation, senescence, and complement activation^98–100^. It is possible that CD8+ T cell and NK cells may produce inflammatory mediators upon weight loss that impact the local environment and affect adipose expansion following weight regain. Future studies should more specifically examine how distinct antigen presentation pathways in the adipose tissue impact specific effector subtypes and their functional state with weight loss and how these changes further affect local and systemic metabolism following weight regain.

There are a few limitations to our study. First, this study was conducted exclusively in male mice, because female mice have attenuated weight gain and metabolic function to high fat diet and attenuated glucose impairments to weight cycling^20^. However, dendritic cells in female mice also have more potent antigen presentation^101,102^, and thus, future research using both male and female mice may help highlight the role of antigen presentation through dendritic cells on weight cycling-mediated outcomes. Another limitation is that we did not assess immune cell populations in adipose tissue from humans with weight loss or weight regain. Recent studies suggest that weight loss via bariatric surgery also induces persistent cellular and transcriptional changes in human adipose tissue^103^, suggesting potential overlap with published murine work (cite Cottam). Additionally, we only assessed protein expression of signal 1 (MHC molecules), signal 2 (costimulatory markers), and signal 3 (TNF) in dendritic cells, rather than performing functional co-culture assays to directly evaluate antigen presentation capacity. However, the high expression of these essential signals required for effective antigen presentation suggests enhanced antigen presenting potential^104,105^. Future research should measure not only functional antigen presentation but the downstream impacts on adaptive cell activation or inhibition and the overall impacts to systemic metabolism, as there is still much to learn about the role of these cells in adipose homeostasis and dysfunction with weight cycling.

In sum, our work suggests that weight loss notably drives dendritic cell MHCIb antigen presentation and the expansion of CD8+ T cell, NKT, and NK populations with concurrent exhaustion. Both non-classical MHCIb antigen-presentation and effector cell exhaustion are underexplored areas of molecular regulation in adipose tissue, and our results open many new questions including which antigens are presented with weight loss. Moreover, this work suggests weight loss modulates multiple dendritic cells subsets, MHC molecules, and effector populations that may affect not only the local adipose tissue, but systemic metabolism following weight regain.

## Acknowledgements

This work has been pre-printed on bioRxiv.

## Funding

This study was supported by a Texas American College of Sports Medicine (TACSM) Student Research Development Award (SRDA) to WK, an American Heart Association (AHA) Transformational Project Award to HC (25TPA1471583) and a Houston-Nutrition and Obesity Research Center Pilot and Feasibility Award to HC.

## Conflicts of Interest

None declared.

## Author Contributions

WK and HLC conceptualized this project. WK, MK, EM, and HLC contributed to the data collection, and WK analyzed the data and wrote the initial draft of this work. WK, ECL, HW, and HLC contributed to data interpretation, and all authors provided comments or edits and approved this work for submission.

## Notes

### Competing Interest Statement

The authors have declared no competing interest.

