## Supplemental Figures for "Weight loss increases adipose dendritic cells, non-classical antigen presenting proteins, and cytotoxic effector cells"

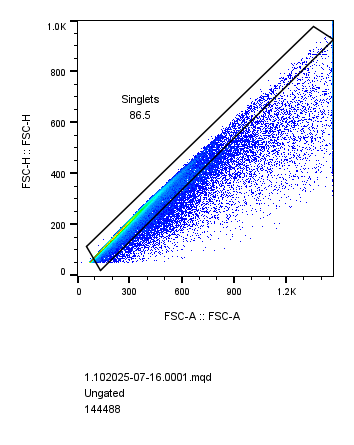

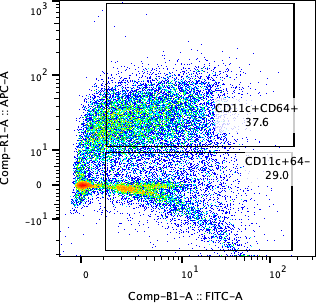

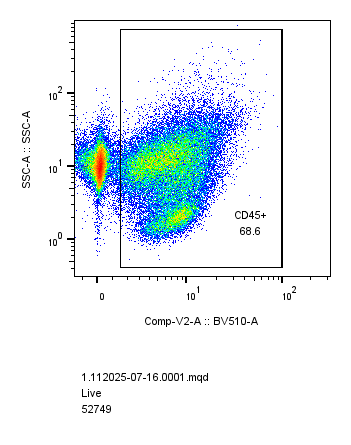

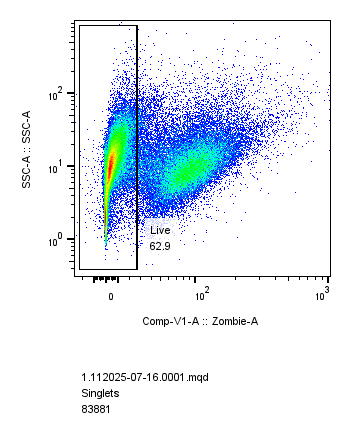
 **A**	Singlets	Live	CD45	DC or ATM

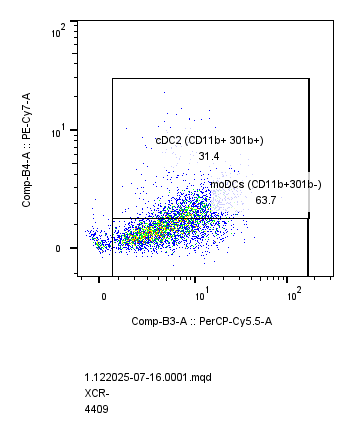

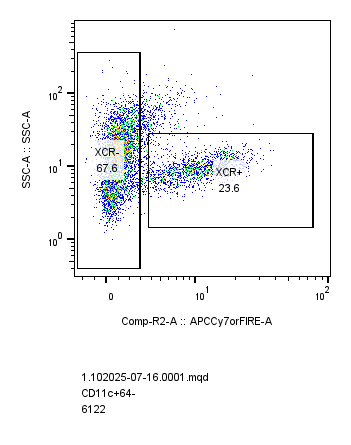
**B**	cDC1	cDC2 and moDC

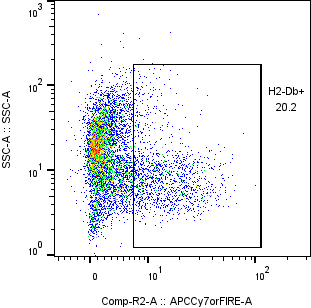

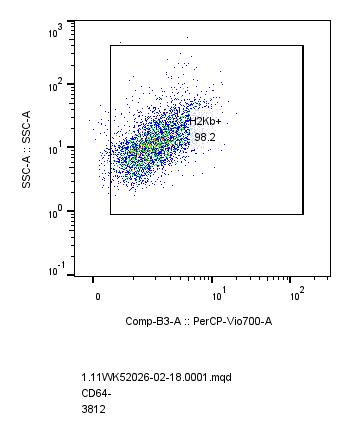

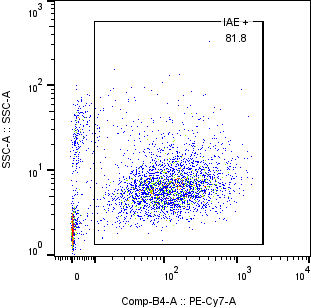
**C**	IA/IE	**D**	H2-Db	H2-Kb

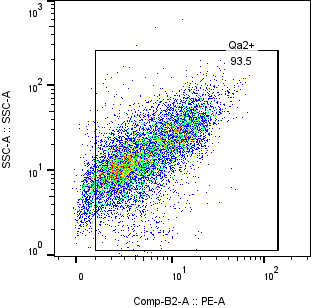

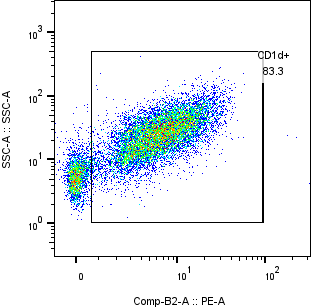

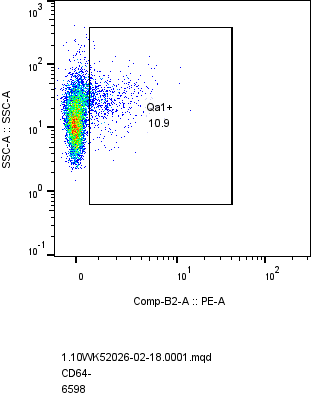
**E**	Qa1	Qa2	Cd1d

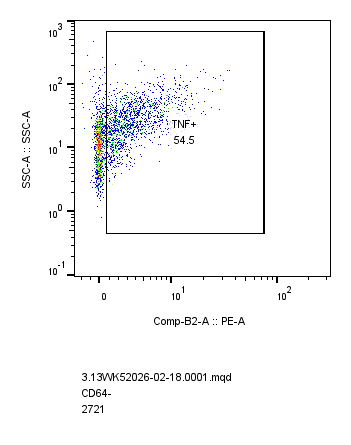

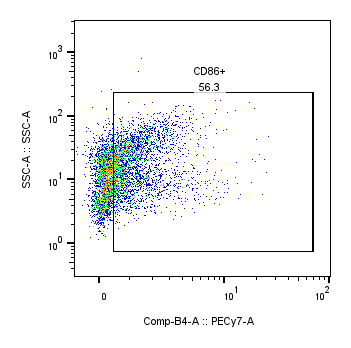

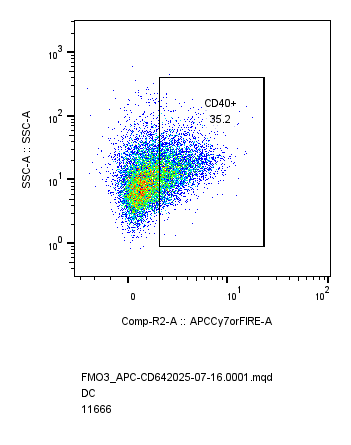
**F**	CD40	CD86	**G**	TNF

**Figure S1: Gating strategy of adipose dendritic and macrophage cells with MHC, Costimulatory and TNF molecules.** A) Cells were gated from singlets, live, CD45+ to, CD11c+64- as dendritic cells and CD11c+64+ as macrophages. B) cDC1 were gated on XCR+ from dendritic cells, cDC2 were gated on XCR-CD11b+301b+ from dendritic cells, and moDCs were gated on XCR- CD11b+301b- from dendritic cells. C) IA/IE, D) H2Db+ and H2Kb, E) Qa1+, Qa2+ and CD1d+, F) CD40+ and CD86+, and G) TNF+ were all gated from dendritic cells.

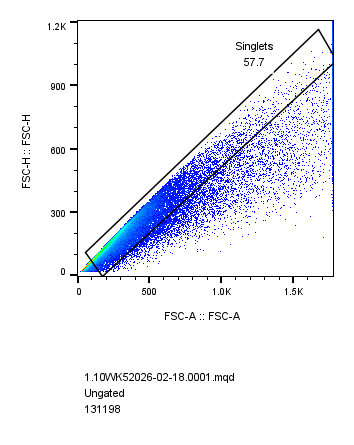

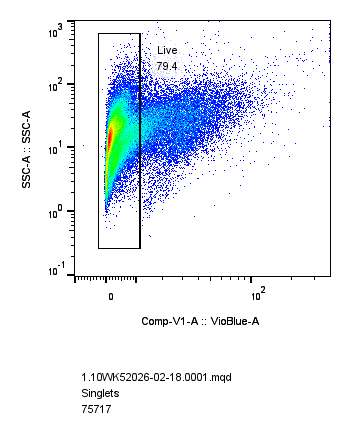

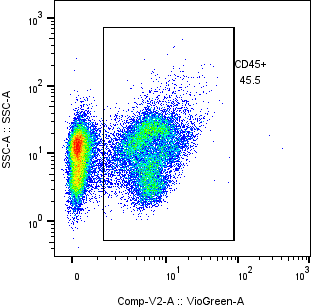
 Singlets	Live	CD45

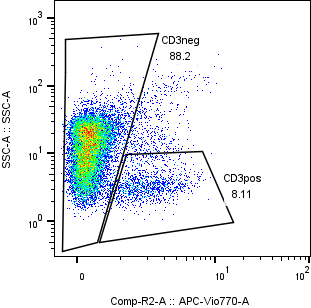

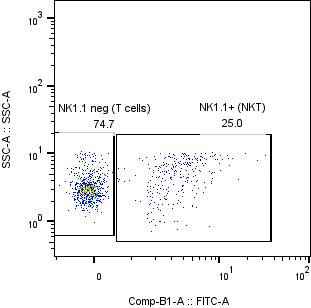

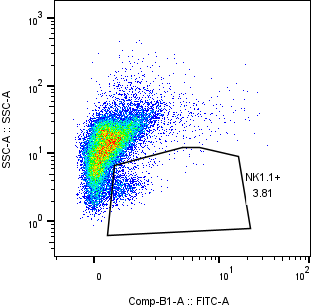
**A**	NK Cells	**B**	NKT cells

Lean

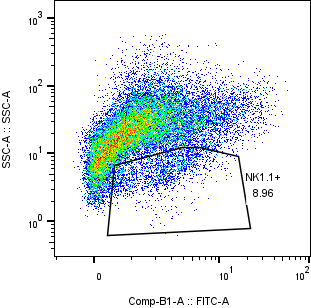

Weight loss

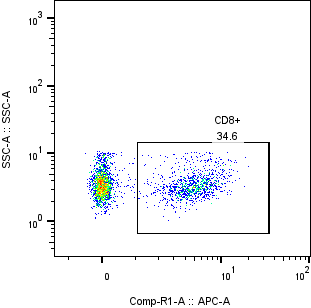
**C**	CD8+ T cells

**Figure S2: Gating strategy adipose NK, NKT and CD8+ T cells**. Cells were gated from singlets, live, CD45+ to A) CD3- NK1.1+ as NK cells, B) CD3+NK1.1+ as NKT cells (lean and weight loss samples shown), and C) CD3+NK1.1- CD8+ as CD8 T cells.

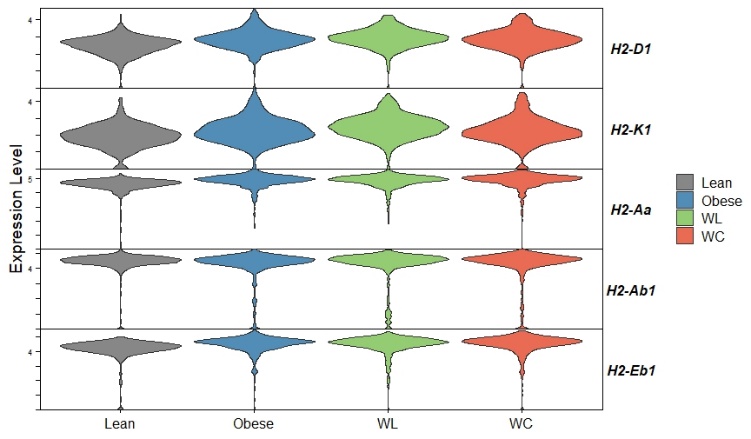

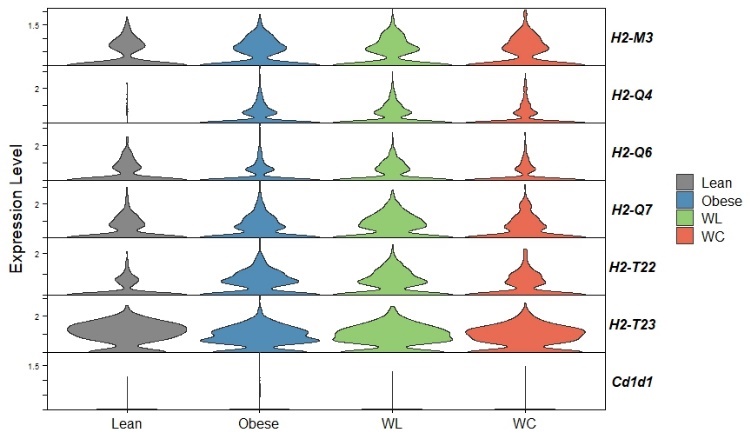
 **A**	cDC1

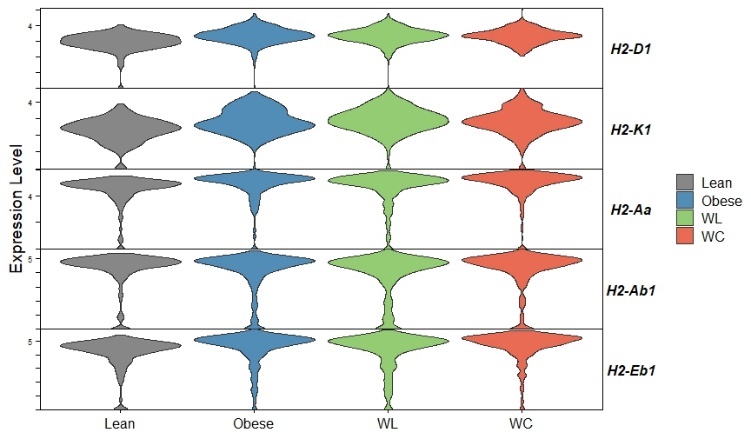

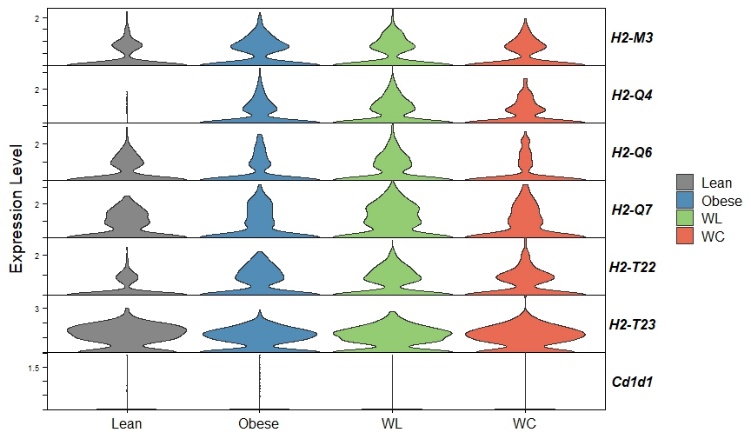
**B**	cDC2

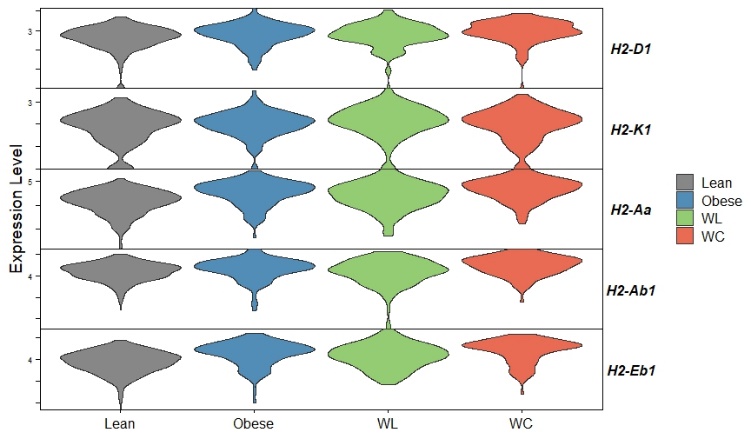

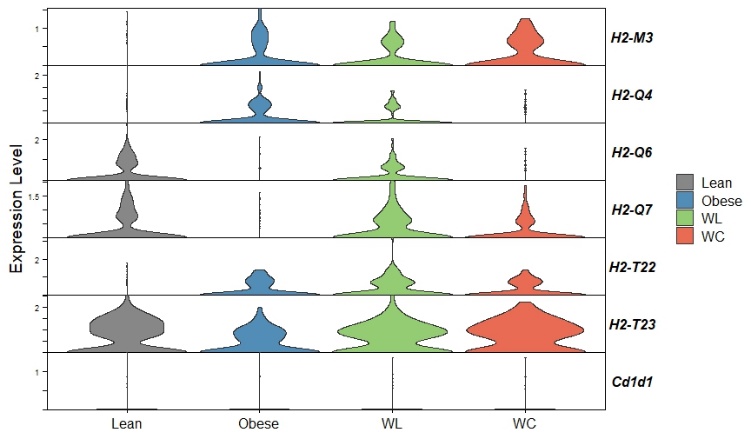
**C**	moDC

**Figure S3: *H2-T23 (*Qa1*)* increases in moDC and *H2-Q6 and H2-Q7 (*Qa2*)* seemingly increase with weight loss in cDC1 and cDC2.** Non-classical MHCIb expression and classical MHCI and MHCII expression in dendritic cell (DC) subsets (A) conventional (c)DC1, (B) cDC2, and (C) monocyte-derived (mo)DC from lean, weight gain (WG), weight loss (WL), and weight cycled (WC) mice from single-cell RNA sequencing analysis of data of Cottam et al 2022.

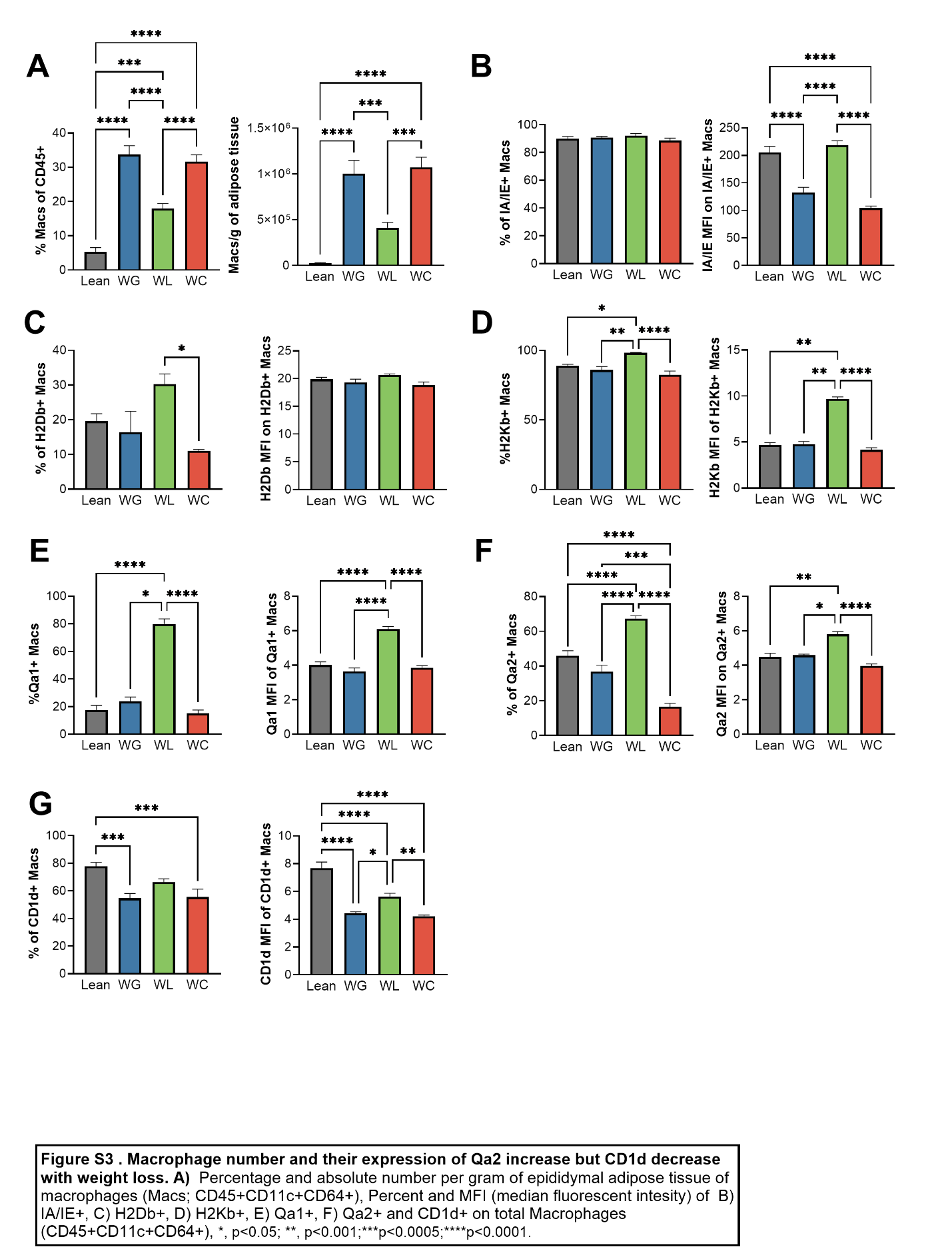

**Figure S4. Macrophage number and their expression of Qa2 increase but CD1d decrease with weight loss.** A) Percentage and absolute number per gram of epididymal adipose tissue of macrophages (Macs; CD45+CD11c+CD64+), and percent and MFI (median fluorescent intensity) of B) IA/IE+, C) H2Db+, D) H2Kb+, E) Qa1+, F) Qa2+ and CD1d+ on total macrophages (CD45+CD11c+CD64+), *, p<0.05; **, p<0.001;***p<0.0005;****p<0.0001.

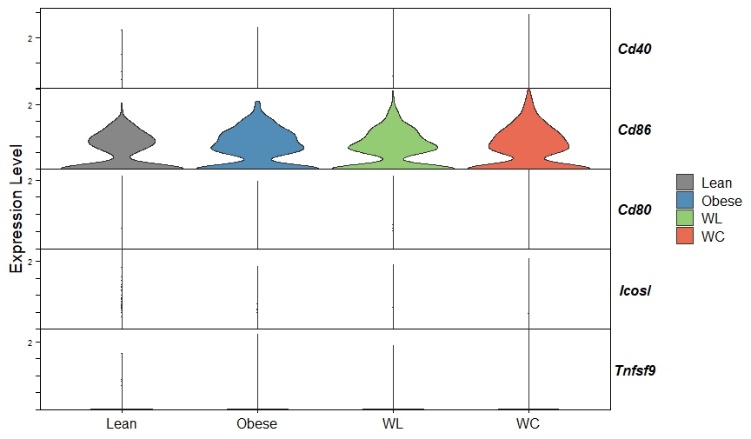
 **A**	cDC1

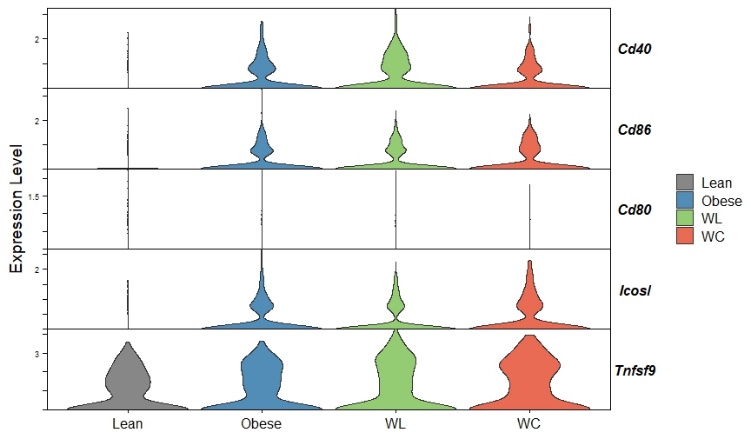
**B**	cDC2

**C**	moDC

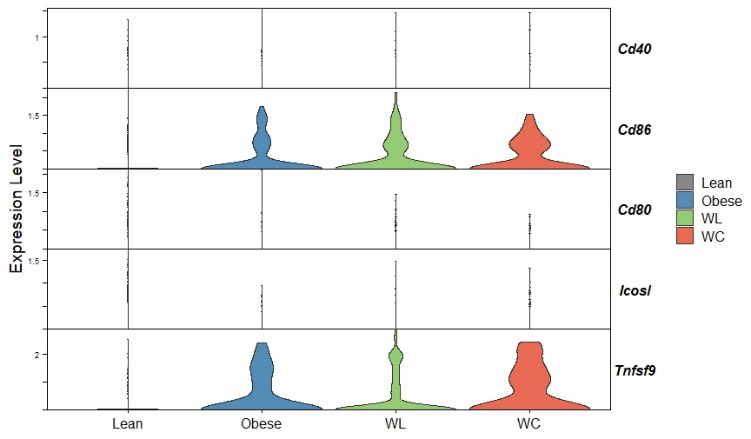

**Figure S5: Expression of *Cd86* increases with weight loss in moDCs and stays elevated following weight gain in cDC1 and cDC2**. Costimulatory molecules in dendritic cell (DC) subsets (A) conventional (c)DC1, (B) cDC2, and (C) monocyte-derived (mo)DC in lean, weight gain (WG), weight loss (WL), and weight cycled (WC) mice from single-cell RNA sequencing analysis of data of Cottam et al 2022.

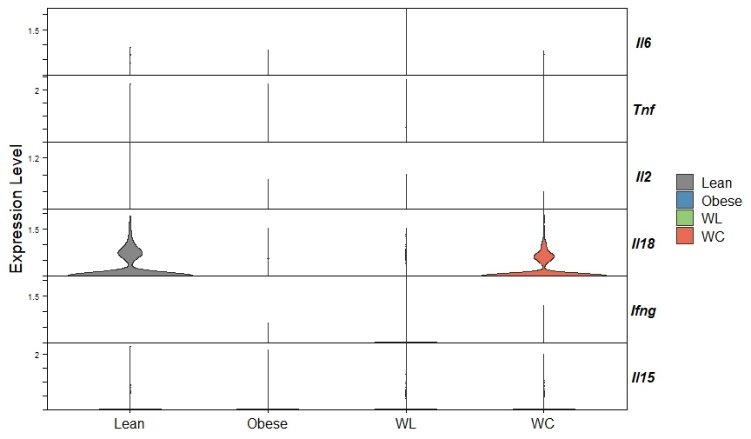
**A**	cDC1

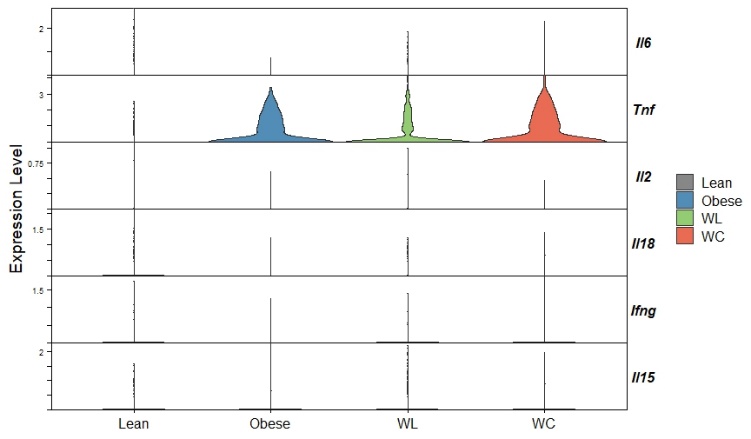
**B**	cDC2

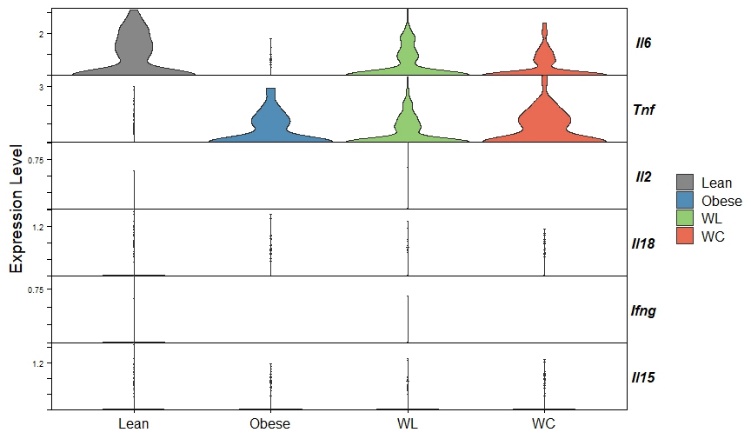
**C**	moDC

**Figure S6: Gene expression of inflammatory cytokines increase differential in each population.** Inflammatory cytokine expression in dendritic cell subsets (A) conventional (c)DC1, (B) cDC2, and (C) monocyte-derived (mo)DC) in lean, weight gain (WG), weight loss (WL), and weight cycled (WC) mice from single-cell RNA sequencing analysis of data of Cottam et al 2022.

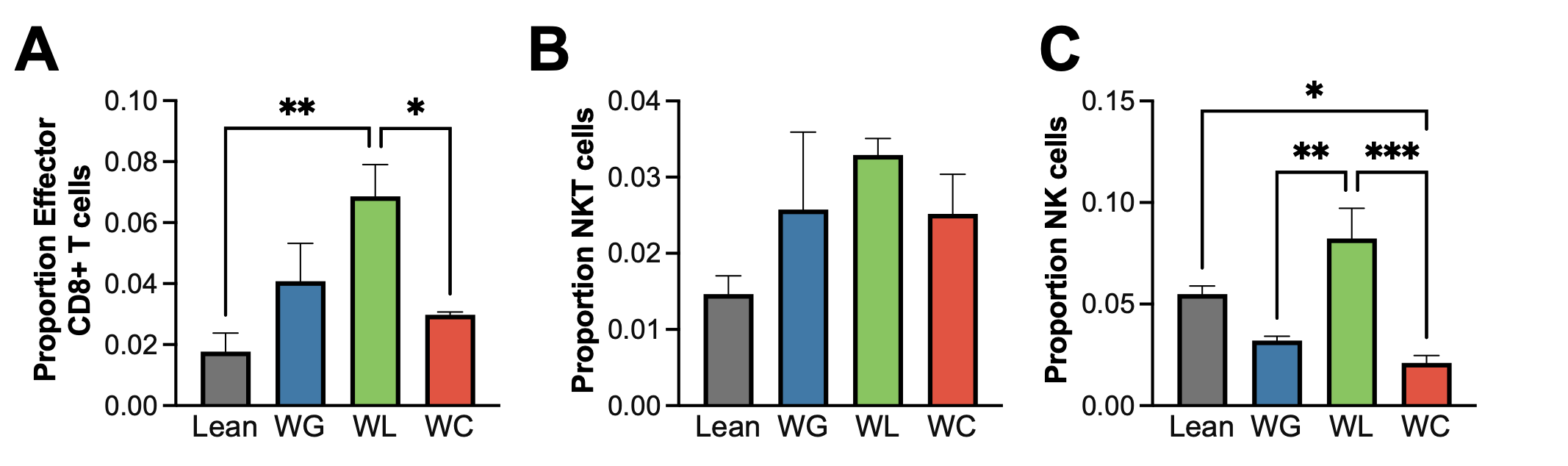

**Figure S7: Effector cytotoxic immune populations increase with weight loss by single RNA-cell sequencing.** (A) Effector memory CD8+ T cells, (B) natural killer (NK)T cells, and (C) NK cells in lean, weight gain (WG), weight loss (WL), and weight cycled (WC) mice from single-cell RNA sequencing analysis of data of Cottam et al 2022.

**A**

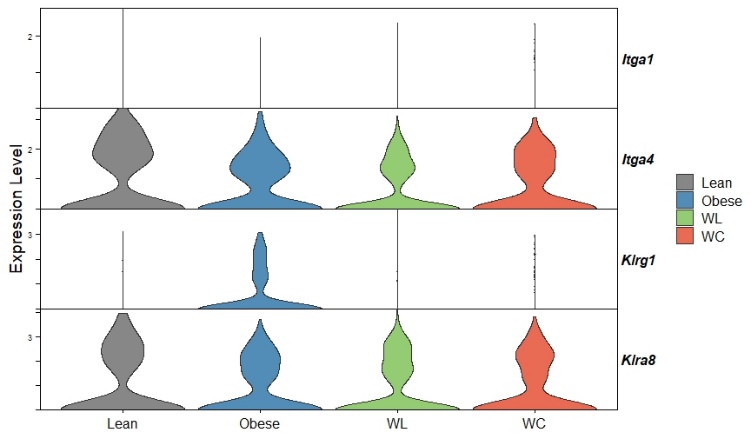

**B**

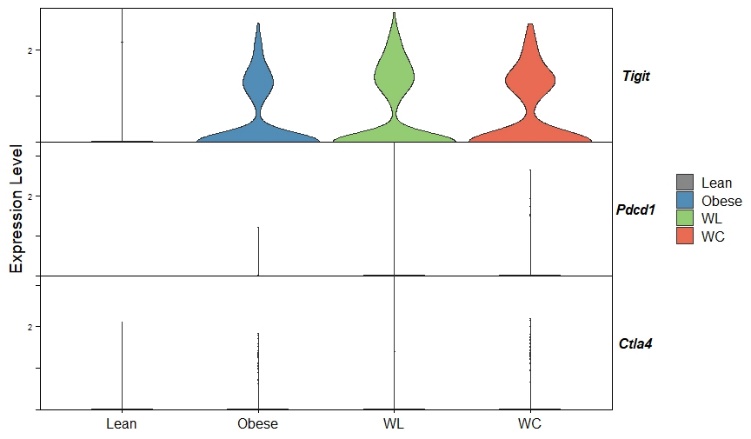

**C**

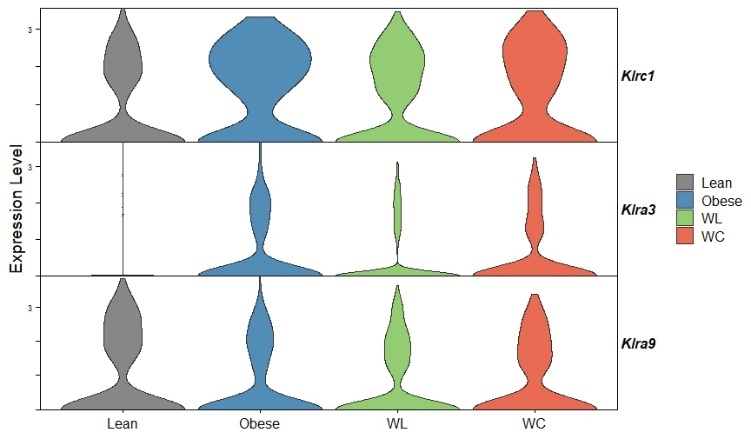

**Figure S8: Genes for activating and inhibitory markers decrease but exhaustive markers increase with weight gain, weight loss and weight cycling in NK cells** (A) Activating, (B) exhaustive (C), and inhibitory markers in lean, weight gain (WG), weight loss (WL), and weight cycled (WC) mice from single-cell RNA sequencing analysis of data of Cottam et al 2022.

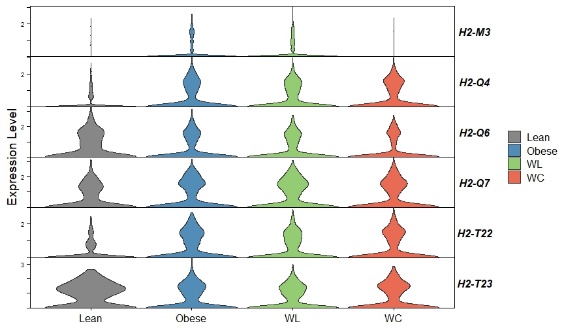

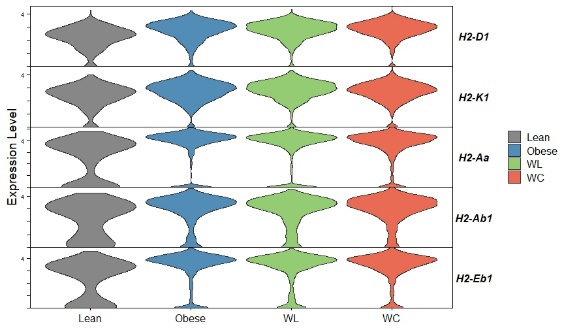

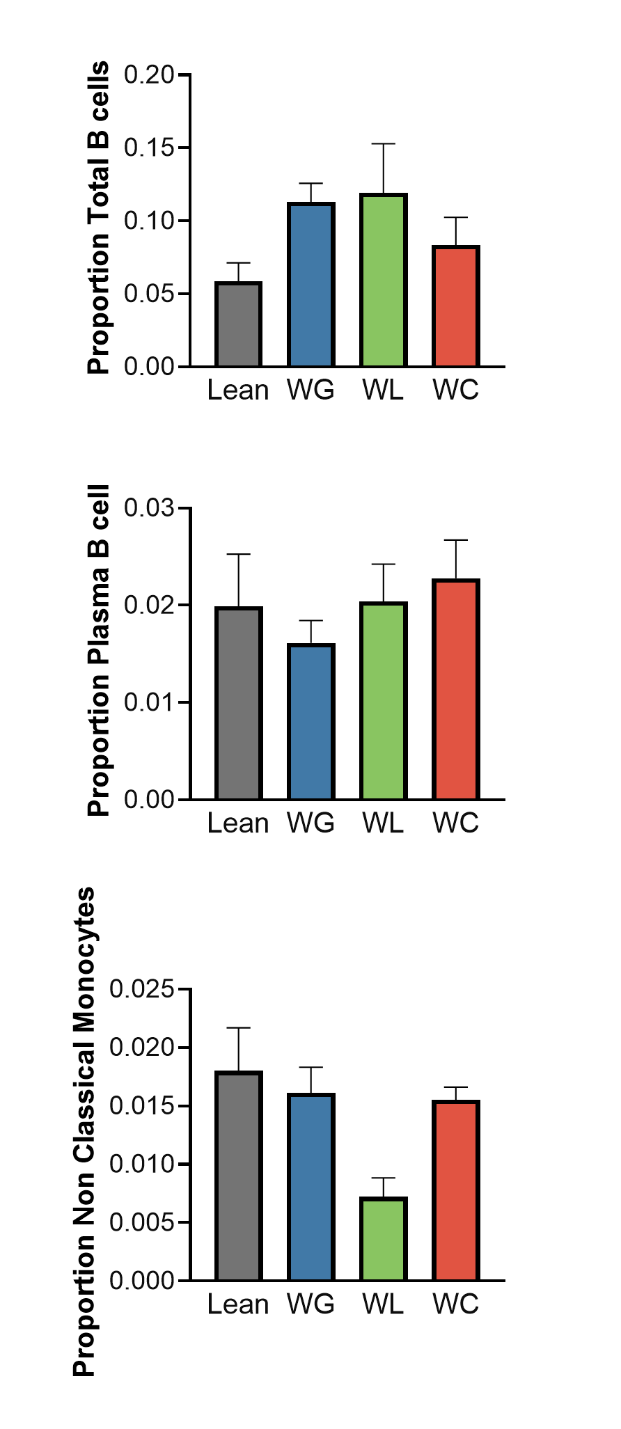
**A**

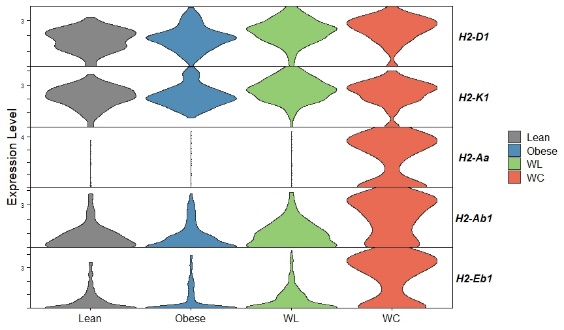

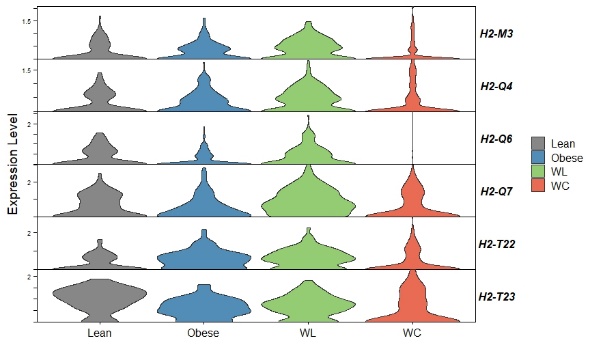

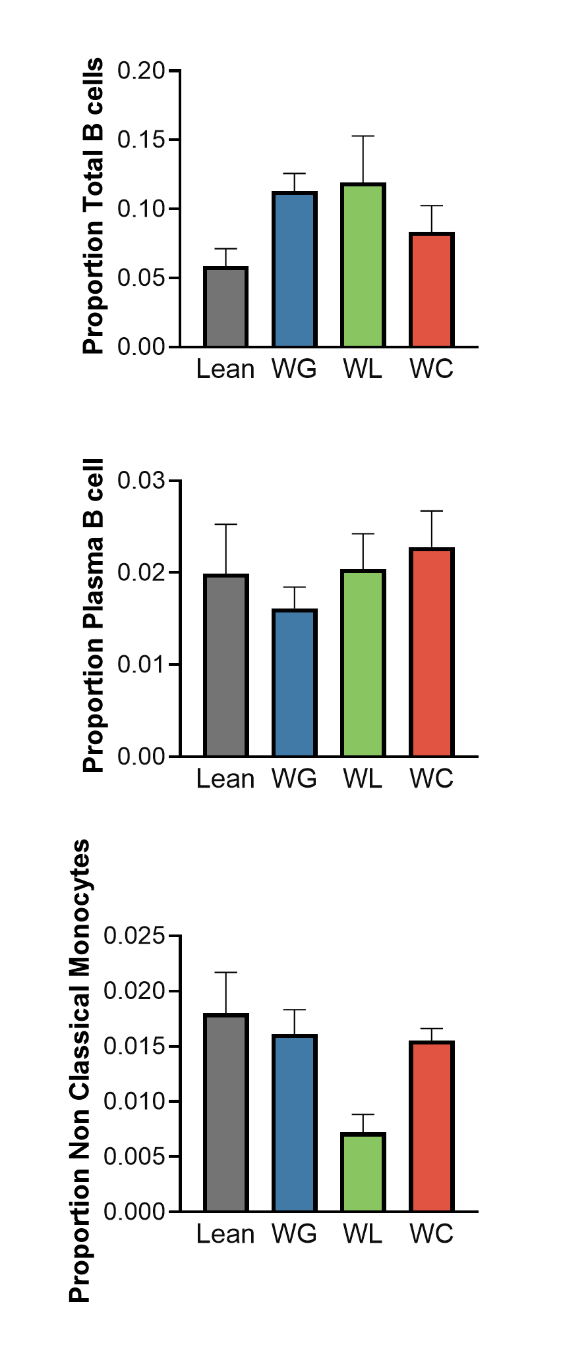
**B**

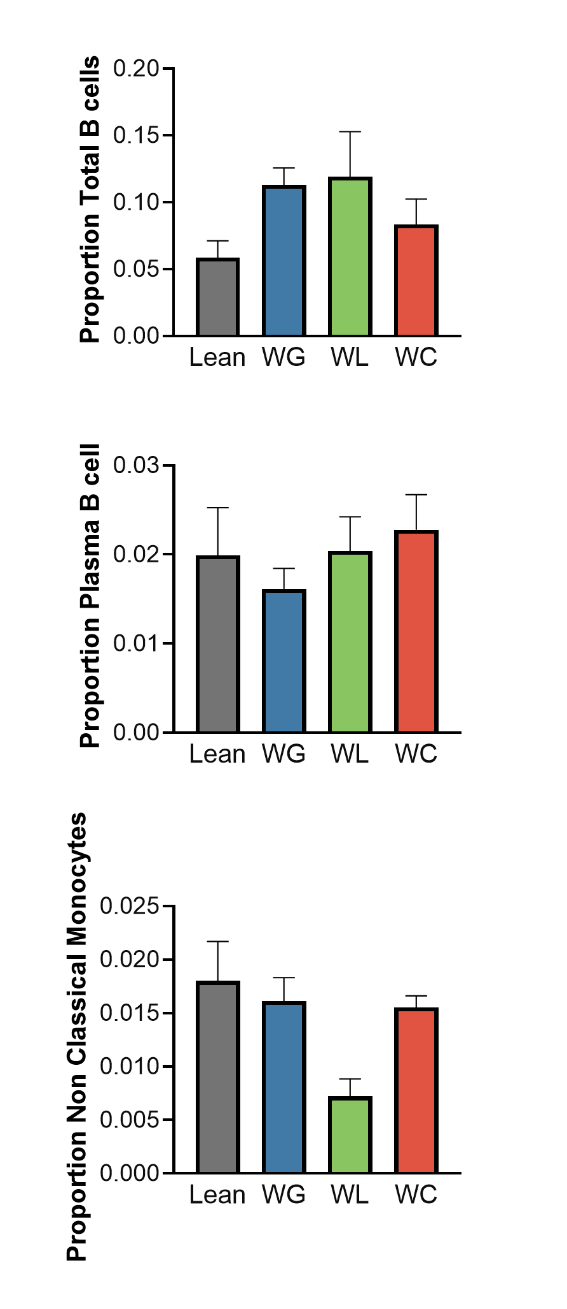
**C**

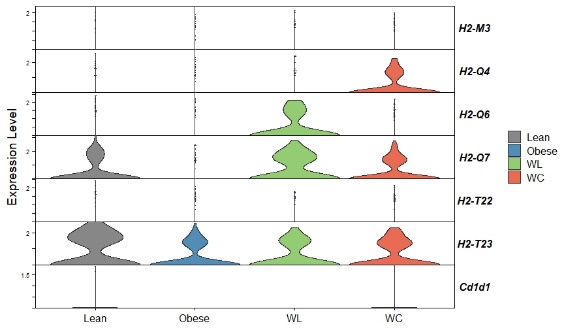

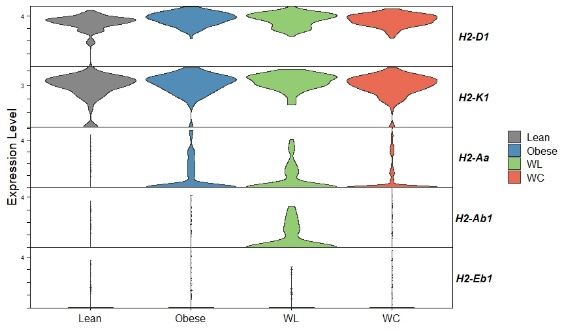

**Figure S9: B cell and non-classical monocytes population with classical and non- classical MHCs.** (A) Total B cells (B) plasma B cells (C) nonclassical monocytes in lean, weight gain (WG), weight loss (WL), and weight cycled (WC) mice from single-cell RNA sequencing analysis of data of Cottam et al 2022.
